# Private Antibody Repertoires Are Public

**DOI:** 10.1101/2020.06.18.159699

**Authors:** Rohit Arora, Ramy Arnaout

**Affiliations:** Division of Clinical Pathology, Department of Pathology, Beth Israel Deaconess Medical Center, Boston, MA 02215.; Division of Clinical Informatics, Department of Medicine, Beth Israel Deaconess Medical Center, Boston, MA 02215.; Harvard Medical School, 25 Shattuck St, Boston, MA 02115

## Abstract

When faced with a given pathogen, the antibody response generally functions similarly across different people,^1–4^ but the source of this similarity has been unclear. One hypothesis was that people share a high proportion of the same VDJ-recombined antibody genes, but this has been disproven.^5,6^ An alternative is that people share a high proportion of *functionally similar* antibodies,^7,8^ but testing this hypothesis requires a method for measuring functional similarity that scales to the millions of antibodies per repertoire and across multiple repertoires, which is impossible experimentally. We recently described a framework for doing so computationally,^9^ which revealed that repertoires consist of loose overlapping functional classes of antibodies with similar antigen-binding capacities;^10–12^ this framework allowed us to estimate a repertoire’s antigen-binding capacity, *τ*, for the ideal target of any given antibody. Here, we show that this framework supports the second hypothesis, and provide the first comprehensive demonstration of overwhelming functional overlap between repertoires from 20 different individuals directly from sequence, without need of binding studies. Overlap is highest among the young and falls with age, due to the selective loss of antibodies that represent a core set of shared or “public” antigen-binding capacities. We reveal considerable heterogeneity in antigen-binding capacities for antibodies against influenza, HIV, and SARS-CoV-2, and show that while some of these classes shrink with age, others persist across individuals. These discoveries change our understanding of repertoire diversity and have implications for vaccine and therapeutic-antibody development, especially for the aged.

---

Recent large-scale studies have shown that <1% of IgG heavy-chain CDR3s are measurably shared across people: the so-called public repertoire of shared sequences is small, dwarfed by the >99% of the repertoire that is “private” or effectively unique to a given individual.^5,6^ The questions arise as to whether public sequences are responsible for public or shared functionality, and what role the much larger private repertoire may play. It is well known that different antibodies can bind a given antigen if they are structurally similar, with the degree of similarity influencing the relative strength of the interaction, as measured by *K*_*d*_ or IC_50_^13–18^ (Fig. 1a-c). Thus even without sharing sequences that encode antigen-specific antibodies at meaningful frequencies (Fig. 1d), repertoires from two individuals can display similar antigen-binding capacities (Fig. 1e) indicating public function from private sequence.^1,19,20^ However, determining whether this observation generalizes to repertoires as a whole has been challenging, since experimentally it is impossible to measure all possible antibody-antigen binding interactions or the pairwise structural similarities among all antibodies for even a single repertoire, much less for multiple repertoires from across different individuals for purposes of comparison. Recently we introduced a computational method for investigating repertoire-wide antigen binding, using thousands of *K*_*d*_ measurements for amino-acid substitutions to establish and validate a mean-behavior model for structural similarity among all pairs of antibodies in a repertoire directly from repertoire sequence.^9^ That work revealed that repertoires are organized into loose overlapping classes of antibodies with similar predicted functionality, and suggested that these classes may be widely shared among repertoires from across individuals, notwithstanding low sequence overlap between repertoires.

**Figure 1.**
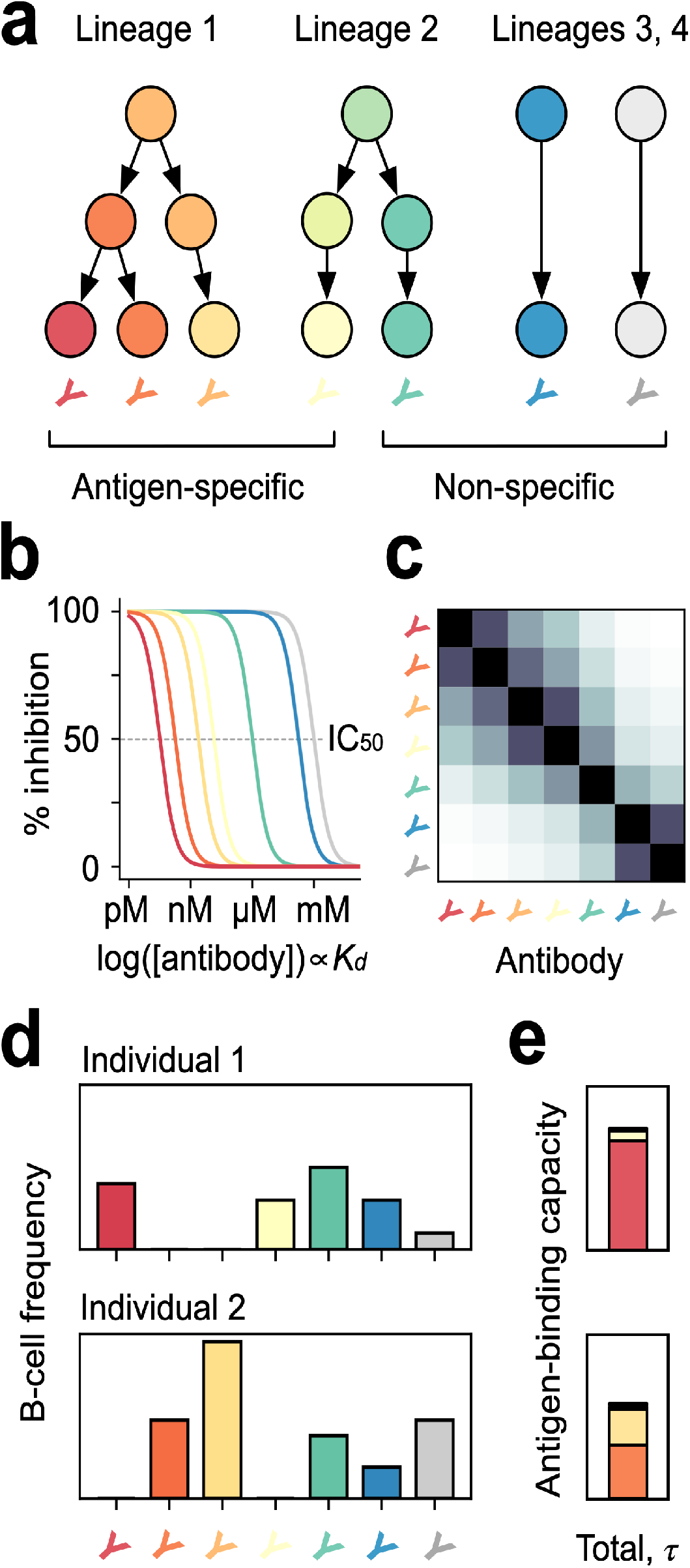
Sequences, classes, and antigen-binding capacity (*τ*). In a repertoire (a), different lineages of B cells can result in antibodies that are considered specific (warm colors; left) or non-specific (cool colors; right) for a given antigen (here the ideal antigen for the red antibody), as measured by *K*_*d*_ or IC_50_ (b). The pairwise similarity between any pair can be quantified^9^ (c); antibodies with similar antigen-binding specificity define a functional class. Multiplying by B-cell frequency (d) and then summing the resulting similarity-weighted frequencies yields ***τ***, the repertoire’s total binding capacity for this antigen (e). Note that the antibody with the highest specificity (red) need not be present in a repertoire, for a repertoire to have substantial antigen-binding capacity against its preferred antigen (d-e, bottom panels).

In the present work, we direct tested this possibility by measuring the overlap of functional classes between IgG repertoires from different individuals. Specifically, we measured how representative^21^ of each other IgG CDR3_H_ repertoires are, using a cohort of 20 individuals, 10 aged 21-27 years and 10 aged 73-93 years, which for convenience we refer to as “young” and “old”.^22^ Representativeness, written as the symbol 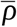 (“rho-bar”), is a mathematical measure of normalized beta (intergroup) diversity^21^ that indicates how typical one repertoire is of a set of repertoires (Methods). For comparing pairs of repertoires, as we do here, 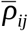 near ½ means repertoire *i* is distinct, overlapping little with repertoire *j*, whereas 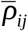 near 1 means repertoire *i* is highly representative of the pair, indicating substantial overlap with *j*. Note that 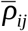 and 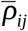 will usually differ; they will be equal only if overlap between them is symmetrical, and will differ most if one repertoire contains the other. 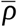 can be scaled and normalized to define an overlap that is 0 at a 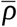 of ½ and 1 at a 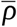 of 1, which can be applied to sequences or functional classes.

We first used 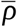 to measure sequence overlap for pairs of young repertoires (Fig. 2a). As expected,^5,6^ overlap was essentially nonexistent (0.00±0.01), reflecting negligible sequence overlap between any pair of these repertoires (range, 5,267-35,811 unique and 10^5^ total sequences per repertoire). In stark contrast, overlap for functional classes was high (0.62±0.08), reflecting substantial overlap between repertoires at a class level (Fig. 2b). This result indicates that public function from private sequence is a dominant feature of IgG repertoires of the young and can be traced to functional similarity encoded in the sequence of CDR3s^9^: antibody repertoires of young individuals overlap almost completely in functional composition, suggesting a large public functional repertoire at the core of the healthy human immune system. Antibody sequence diversity generally falls with age.^23,24^ We recently showed the same is true for functional classes in T-cell receptor repertoires, with some intriguing exceptions.^9^ We next asked what effect a fall in IgG CDR3_H_ sequence diversity might have on functional overlap between individuals. As with the young repertoires, we first measured overlap on sequences for all pairs of old repertoires, and as with the young repertoires, we found very low sequence overlap (0.04±0.12) (Fig. 2c), although it was higher in old-old pairs than young-young pairs (Mann-Whitney U [MWU] *p*=8.9×10^−6^) (Fig. 2e).

**Figure 2.**
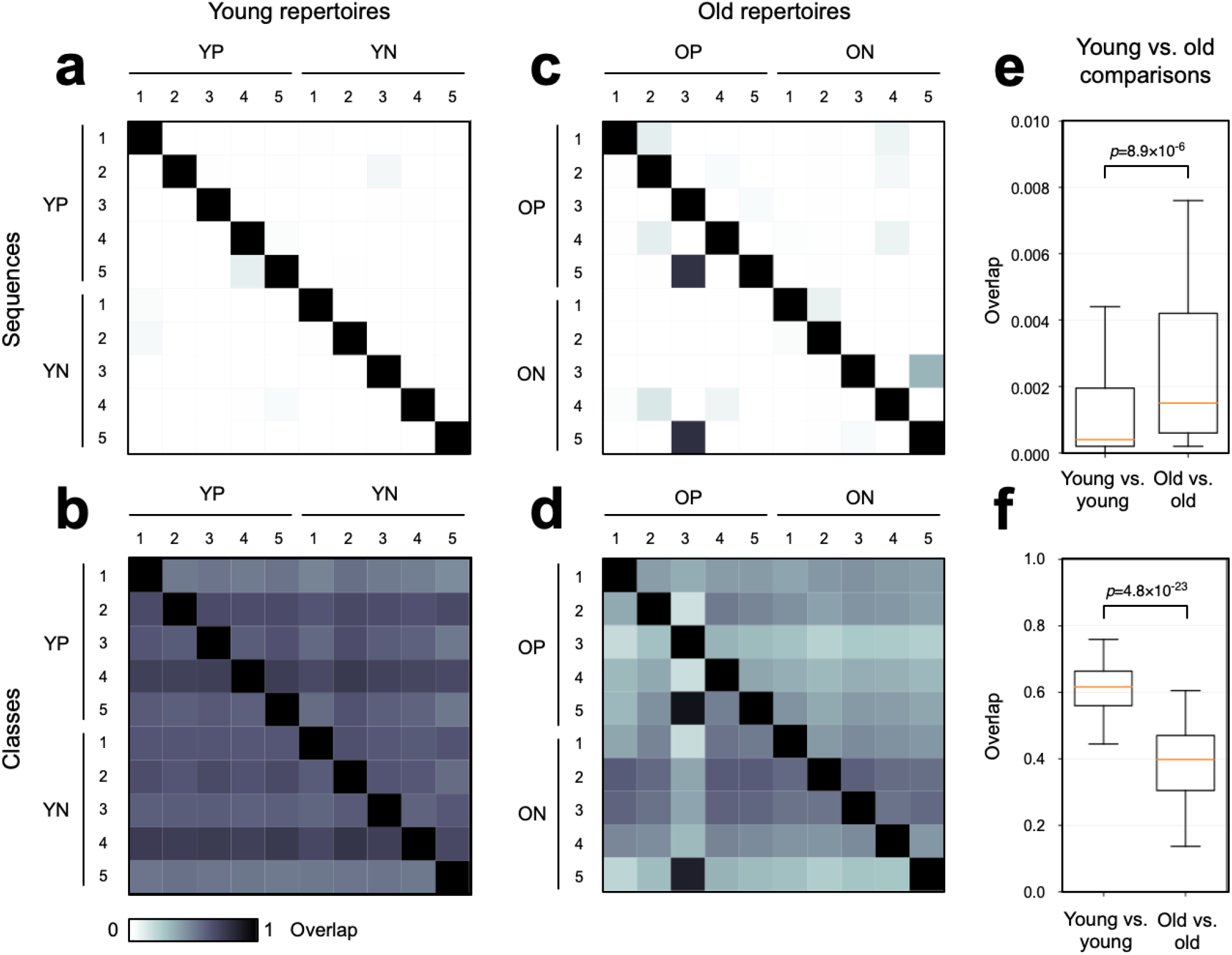
Sequence and class overlap between repertoires. Sequence (a) and class (b) overlap between pairs of young repertoires, grouped by CMV serostatus (YP, positive; YN, negative), and sequence (c) and class (d) overlap between pairs of old repertoires, again grouped by CMV serostatus (OP, positive; ON, negative). Box-and-whiskers plots comparing overlap between young-young pairs and old-old pairs for sequence (e) and class (f) overlap.

Again in contrast, overlap for functional classes (Fig. 2d) was much higher than it was for sequence (0.40±0.14) but overlap for the old was also substantially lower than it was for functional classes between pairs of young individuals (MWU *p*=4.8×10^−23^) (Fig. 2f). There were no obvious differences by cytomegalovirus (CMV) serostatus. Thus, aging diversifies antibody repertoires not only away from their common beginnings, but also away from each other.

We next sought to characterize how old repertoires diversify, to determine whether for example diversification reflects a wholesale replacement of functionalities present in young repertoires, by others specific to each individual as they age. Replacement would result in 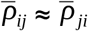 for a given pair of young-and-old repertoires, i.e. symmetric overlap: each repertoire would be similarly representative of the pair. We found that the overlap was moderately directional: young IgG repertoires contain much of the functionality old ones, with 0.56±0.11 for overlap of young with old (i.e., based on how representative the young representative is of the pair) vs. 0.43±0.11 for overlap of old with young (Fig. S1a-b). Thus, aging of the IgG repertoire is more a process of expansion of existing functionalities than invention of new ones^25–27^, a “privatization” marked more by selection and/or drift than gain of new functionality.

We next asked whether any functionalities increase or decrease in common across individuals, perhaps in response to common infectious exposures,^28,29^ or whether differences are on the whole idiosyncratic to each individual. To answer this question, we calculated the total antigen-binding capacity, ***τ*** (Fig. 1e), of each CDR3 for its antibody’s ideal target antigen, for all 1,25,292 sequences present in each of the 10 old repertoires. A sequence with high ***τ*** represents a large functional class.^9^ The class will be large if (*i*) the B cell expressing this sequence is present at high frequency, (*ii*) the repertoire contains many similar sequences, (*iii*) B cells expressing similar sequences are present at high frequencies, or (*iv*) some combination of (*i*)-(*iii*) (Fig. 1d). Note that a sequence can be used to measure ***τ*** in a repertoire that does contain the sequence; for example, the red antibody in Fig. 1d is absent from repertoire 2 but ***τ***_red in repertoire 2 is not zero (it is roughly half of ***τ***_red in repertoire 1): repertoire 2 contains antibodies that can bind the red antibody’s ideal antigen (Fig. 1e).

For each sequence in the old repertoires we calculated its ***τ*** in each of the old repertoires and plotted the average against its average ***τ*** across young repertoires (Figs. 3a-b). Sequences with no change in binding capacity between young and old lie on the diagonal; sequences that represent functionalities that decrease with age lie above the diagonal, while sequences that represent functionalities that increase lie below the diagonal. For this analysis, we excluded the small number of sequences with very high variance in order to identify functionalities that were expanded across all old repertoires (Fig. S3). We observed an excess of sequences above the diagonal that had higher ***τ*** (Fig. 3a), supporting the conclusion of widespread decreased functionality in the old, especially for functionalities with higher antigen-binding capacities. In contrast, functionalities that were expanded across all old repertoires were disproportionately of lower antigen-binding capacity (lower ***τ***; nearer the origin): an “old core” of consistently increased but nonetheless relatively weak public functionalities.

**Figure 3.**
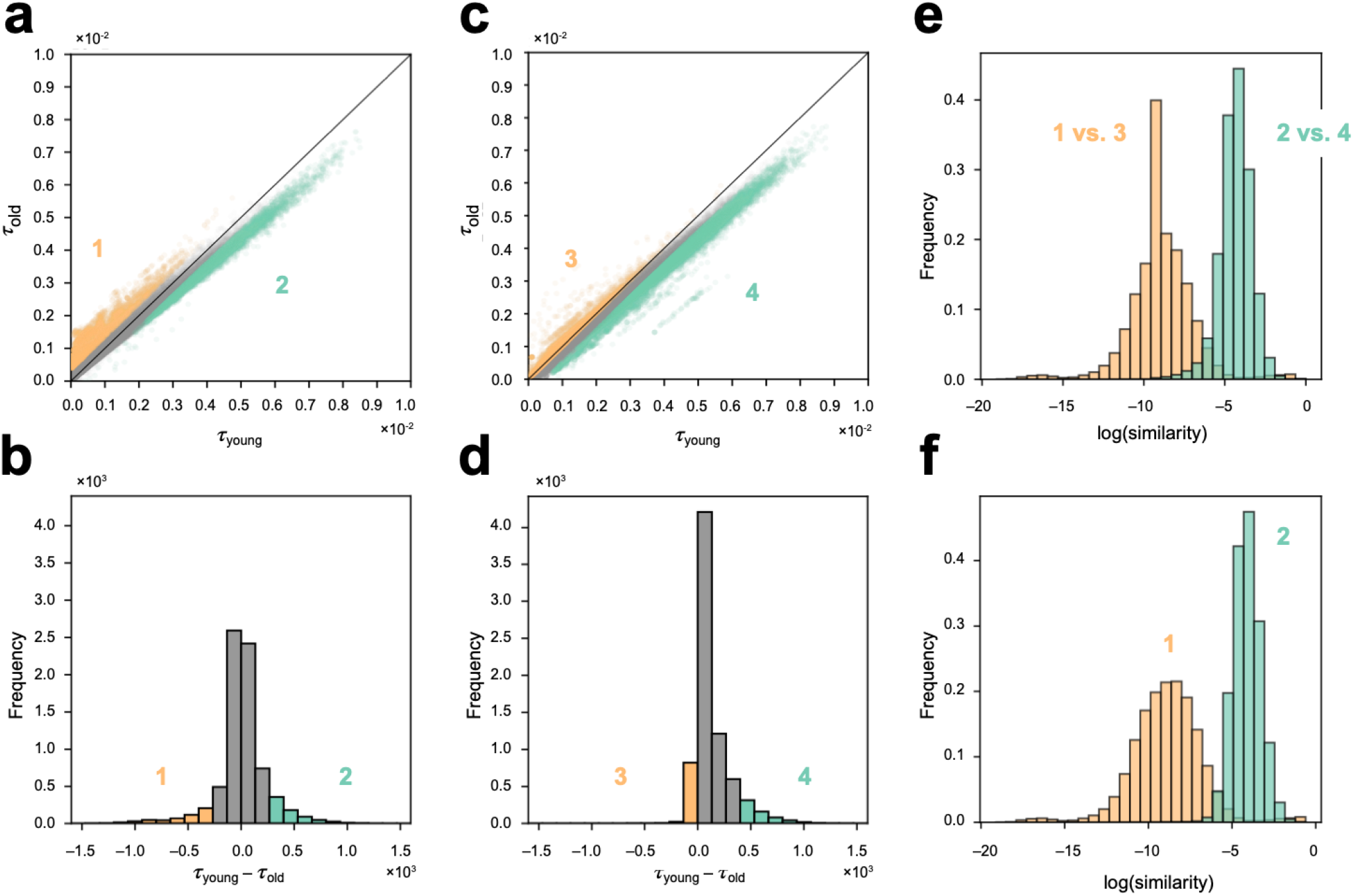
*τ* in young vs. old. Each point represents a single sequence. In (a) all sequences from young repertoires are shown, excluding those with the highest variance in antigen-binding capacity (see Methods); (b) shows the histogram of ***τ***_young_-***τ***_old_ for the sequences in (a). (c) and (d) are analogous but for all sequences from old repertoires, with the same exclusion. In all panels, the 5% with the greatest ***τ***_young_-***τ***_old_ are colored green (the young core), and the lowest 5% are colored yellow (the old core). (e), inter-group similarities between each pair of old-core sequences from (a) vs. (c) (populations 1 vs. 3), and between each pair of young-core sequences from (a) vs. (c) (populations 2 vs. 4). (f), intra-group similarities of sequences belonging to the young core and old core.

We next made the analogous plot for each sequence from the young repertoires (Fig. 3c-d), again with sequences representing functionalities that decrease with age across all old repertoires lying below the diagonal, and those representing increased functionality lying above it. In principle, the four colored regions in Figs. 3a-b and 3c-d represent the same two sets of functionalities, measured in Figs. 3a-b by sequences from old repertoires and in Figs. 3c-d by sequences from young repertoires. To test this prediction, we compared pairwise sequence similarities^9^ (e.g. Fig. 1c) from the colored regions across the plots (Fig. 3e-f). This comparison demonstrated consistency within each set of functionalities and a major difference between them, supporting the view that the plots reflect a diverse old core of relatively weak functionalities that grows with age and a less diverse young core of relatively strong functionalities that decrease with age across individuals (sequences are listed in Supplementary File 1). This result reaffirms the ability of this approach to identify repertoire-wide patterns despite there being almost no public (shared) sequences between young and old (Fig. S1c-d).

We next sought to characterize the antibody sequences that represent these young and old cores. Previous reports have found that IgG CDR3_H_s from repertoires from older individuals are on average slightly longer than those from younger individuals,^18^ and this was technically true in our analysis: 19.1±4.5 residues overall across the young repertoires vs. 18.8±4.4 across the old repertoires, for an average length difference of 0.3 residues, but the histograms exhibited considerable overlap (Fig. 4a). However, the length difference between the cores was over 10 times greater: 19.1±3.8 residues in the old core vs. 13.3±1.7 residues in the young core (Fig. 4b), again demonstrating the differences that classes reveal, relative to analysis of sequence overlap. To begin to characterize other structural differences between antibodies from young and old cores, we mapped each sequence according to its biophysical properties^30^ (excluding length), clustered sequences using UMAP,^31^ and then colored them by core (Fig. 4c). Sequences from the young core were organized into a few large clusters in close proximity to each other, while old-core sequence were organized into many small clusters that were widely separated. This result suggests that the functionalities that define the young core are structurally related, and as these decrease with age they are replaced by a suite of disparate functionalities that are nonetheless broadly shared across individuals.

**Figure 4.**
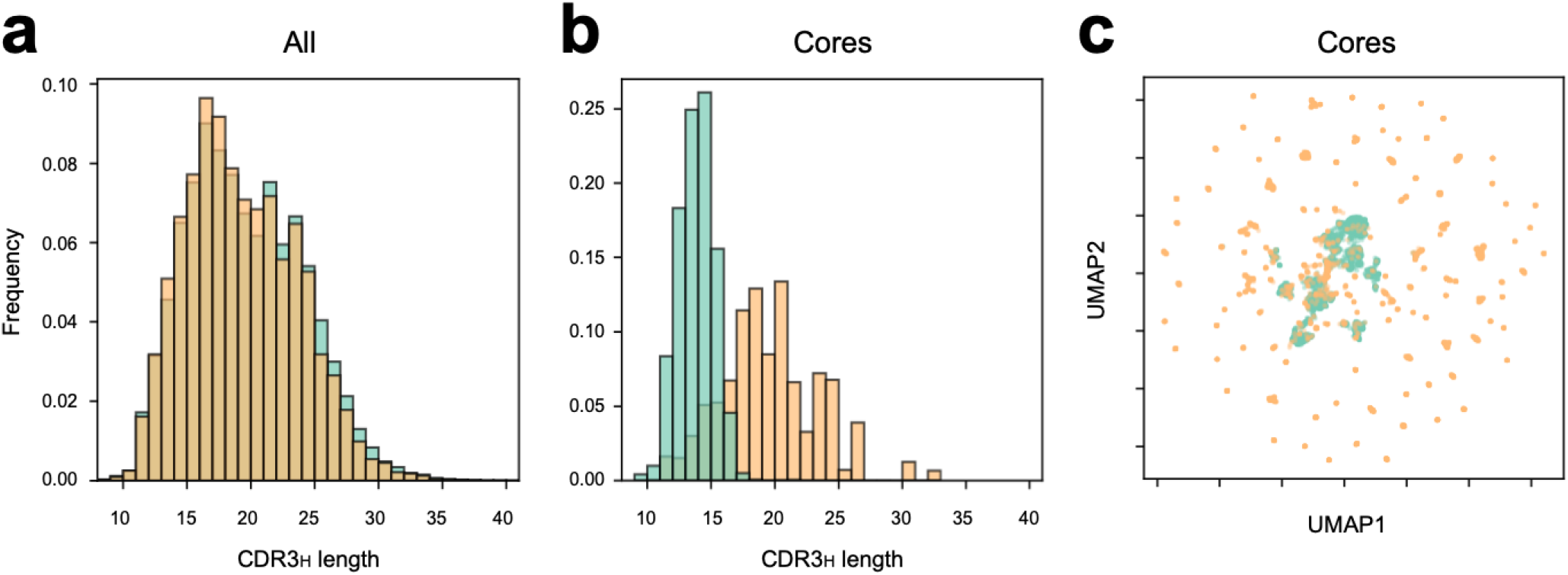
Systematic differences with age. CDR3_H_ length differences between young (green) and old (yellow) repertoires (a) and between young and old cores (b). Clustering young and old cores by biophysical properties.

Core sequences are natural targets for future structural analysis, but the present framework can already be used to ask how the total binding capacity of sequences with known structures might change with age. To illustrate, we extracted CDR3_H_s from 31 HIV gp41/gp120- and 47 influenza A hemagglutinin-specific antibodies and 4 with SARS-CoV and/or SARS-CoV-2-binding capacity present in the Protein Data Bank^29^ (Supplementary File 2), and for each of them, we measured the total antigen-binding capacity (***τ***) in each repertoire and calculated the mean and variance across young repertoires and separately across old repertoires. By doing so, the present framework allows identification of antibodies that represent the highest antigen-binding capacity across the most people, in young or old. We found ***τ*** varied over a wide range, with higher variance in the old than the young, and identified antibodies that represent high antigen-binding capacities and low variance (Fig. 5). This method may prove useful for prioritizing antibodies or antibody responses for therapeutic or vaccine-related uses.

**Figure 5.**
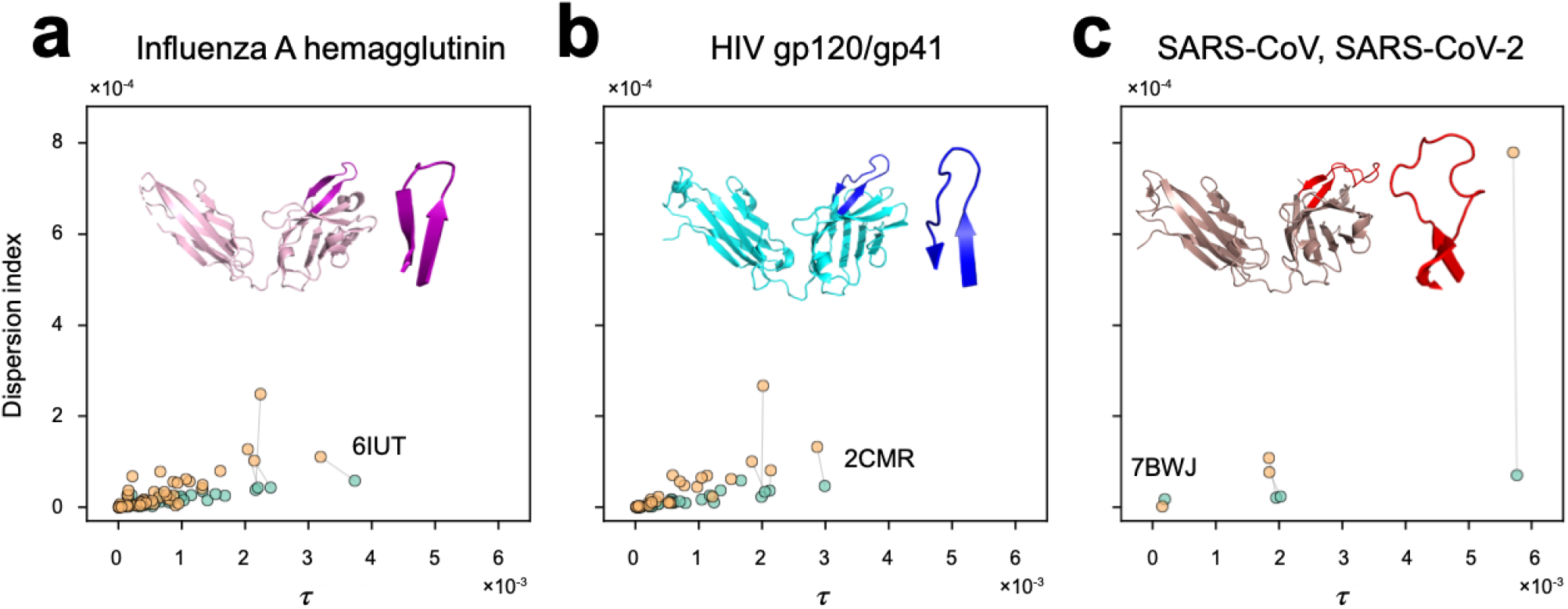
*τ* for antibodies of known specificity. (a) Anti-influenza, (b) anti-HIV, and (c) anti-SARS-CoV/SARS-CoV-2 antibodies obtained from the Protein Data Bank (PDB).^36^ In each set, a gray line connects ***τ***_young_ (green) with ***τ***_old_ (yellow) for the four antibodies with the highest *τ*_young_. Insets, PDB and CDR3_H_ structures for the labeled antibody.

In summary, by considering classes instead of sequences, we demonstrate that IgG mRNA repertoires overlap by as much as three quarters (Fig. 2f). Classes are largely public, in stark contrast to sequence repertoires, which are generally >99% private (Fig. 2e). From a functional/class perspective, IgG repertoires start similar but diverge with age, as individuals lose a limited core of high-binding-capacity functionalities (the young core) in favor of a diverse public suite of other functionalities (the old core) as well as other functionalities private to each individual. It is remarkable to see this result for IgG repertoires, for which somatic hypermutation introduce considerable heterogeneity.^32,33^ We predict class overlap for IgM repertoires will be even higher. Repertoires obtained from DNA are known to show higher class diversity^9^ since they capture sequence from quiescent B cells; it will be interesting to see whether these classes overlap more or less than what we have observed here from mRNA. Currently, the scale of antibody repertoires necessitates using predictive measures of functional pairwise antibody similarity, such as we validated previously,^9^ as the basis for defining classes. However, current measures can almost surely be improved. It will be interesting to test different measures (by swapping out the similarity matrix,^9^ both predictive^20,34^ and one day experimental, in the continuing effort to define the commonalities that emerge from careful analysis of repertoire diversity.

## Methods

See Supplementary online methods.

