## Supplemental methods and figures for "Private Antibody Repertoires Are Public"

**Supplementary online methods**

**High-throughput repertoires.** Quantitative high-throughput IgG repertoires from 10 males aged 21-27 (the “young” cohort; “young repertoires”) and 10 males aged 73-93 (the “old” cohort; “old repertoires”) are as previously described.[1] Subjects presented for influenza vaccination. Repertoires used for the present studies were from before actual administration (i.e., “day 0”). Repertoires were from mRNA of peripheral blood. Cytomegalovirus (CMV) serostatus was available. Repertoires were downloaded, processed and converted to fasta format using NCBI's SRA toolkit.[2] Isotype assignment and CDR3 annotation were performed using IMGT[3] and our in-house pipeline as previously reported.[4]

**Sequence diversity, class diversity, and repertoire overlap.** $10^5$ total sequences were sampled from the sequence-frequency distribution from each repertoire. In repertoires which did not have at least $10^5$ total sequences, the sequences were oversampled according to sequence-frequency distribution for a total of $10^5$ sequences. Sequence and class diversity for $CDR3_H$ were measured as previously described for $q$=0.[5] Overlap was measured by normalizing representativeness[6] from its range of 0.5-to-1 (0.5 is the minimum-possible value for a pair of repertoires) to 0-to-1. To confirm that oversampling from some repertoires did not affect results, a second test was performed in which each repertoire was downsampled to the size of the smallest repertoire (9311 total sequences); as expected, this resulted in no meaningful change the described results.

**Total binding capacity, $\boldsymbol{\tau}$.** The $\boldsymbol{\tau}_i$ of a $CDR3_H$ sequence $\sigma_i$ (Fig. 1e) is the weighted sum of the frequencies of antibodies $\sigma_j$ in the repertoire (Fig. 1d) weighted by the antigen-binding similarity between $i$ and $j$ (Fig. 1c). Note $\sigma_i$ need not be present in a given repertoire to calculate $\boldsymbol{\tau}_i$ (Fig 1d). $\boldsymbol{\tau}$ was measured for every sequence in every repertoire, and the mean and dispersion index (= variance/ mean) was calculated over all young repertoires and separately over all old repertoires. The repertoire for subject EP3 was found to be dominated by a few very large clones as seen previously in leukemia[7,8] and therefore considered an outlier and not included in calculation of mean and variance. To eliminate outlier sequences in old individuals (that are highly overrepresented in a single old individual) in order to focus on cores, the maximum variance of old sequences across young repertoires (rounded to the nearest millionth) was used as a cutoff and any sequences with variance higher than the cutoff were eliminated; this resulted in filtering out of only ~2% of sequences in the old repertoires and ~0.004% of sequences in the young repertoires (Figure S2). From the remaining ~98% and ~99.96%, the 5% of sequences with highest $\boldsymbol{\tau}$ across old repertoires, and 5% of sequences with highest $\boldsymbol{\tau}$ across young repertoires were designated as core sequences and analyzed in the main text.

**Biophysical features.** Features for each antibody were calculated as previously described[9] and visualized via nonlinear dimensionality reduction using UMAP.[10]

**Antibodies of known specificity.** Sequences and 3D structures of 31 anti-HIV and 47 anti-influenza antibodies representing multiple epitopes were obtained from the Protein Data Bank[11] and their IgH CDR3s were identified as described above. For each of these sequences the average $\tau$ and dispersion index were calculated as above. 3D rendering was performed using PyMol.[12]

### Supplementary Figure Legends

**Fig. S1. Sequence and class overlap between young and old repertoires.** Class (a) and sequence (c) overlap of young with old repertoires, and class (b) and sequence (d) overlap of old with young repertoires, both grouped by CMV serostatus (OP, old positive; ON, old negative; YP, young positive; YN, young negative). Schematic (e) representing the diversification of aging antibody repertoires away from their common beginnings, and away from each other while retaining a common core of sequences.

**Fig. S2. Identifying variance cutoff for $\tau$.** Variance and mean of $\tau$ for sequences from old repertoires across young (a) and old (b) repertoires, and from young repertoires across young (c) and old (d) repertoires. Black dotted lines represents the cutoff for old (variance = $1.6\times10^{-5}$) and young (variance = $4.1\times10^{-5}$) sequences along the variance axis, beyond which the sequences were eliminated from the analysis.

### Supplementary Files

**File 1.** Microsoft excel format (.xlsx) file containing the list of young and old core IgG $CDR3_H$ sequences. Download here.

**File 2.** Microsoft excel format (.xlsx) file containing the list of specific $CDR3_H$ sequences (anti-HIV, anti-influenza A, anti-SARS-Cov, anti-SARS-Cov2), extracted from PDB. Download here.

### Figure S1

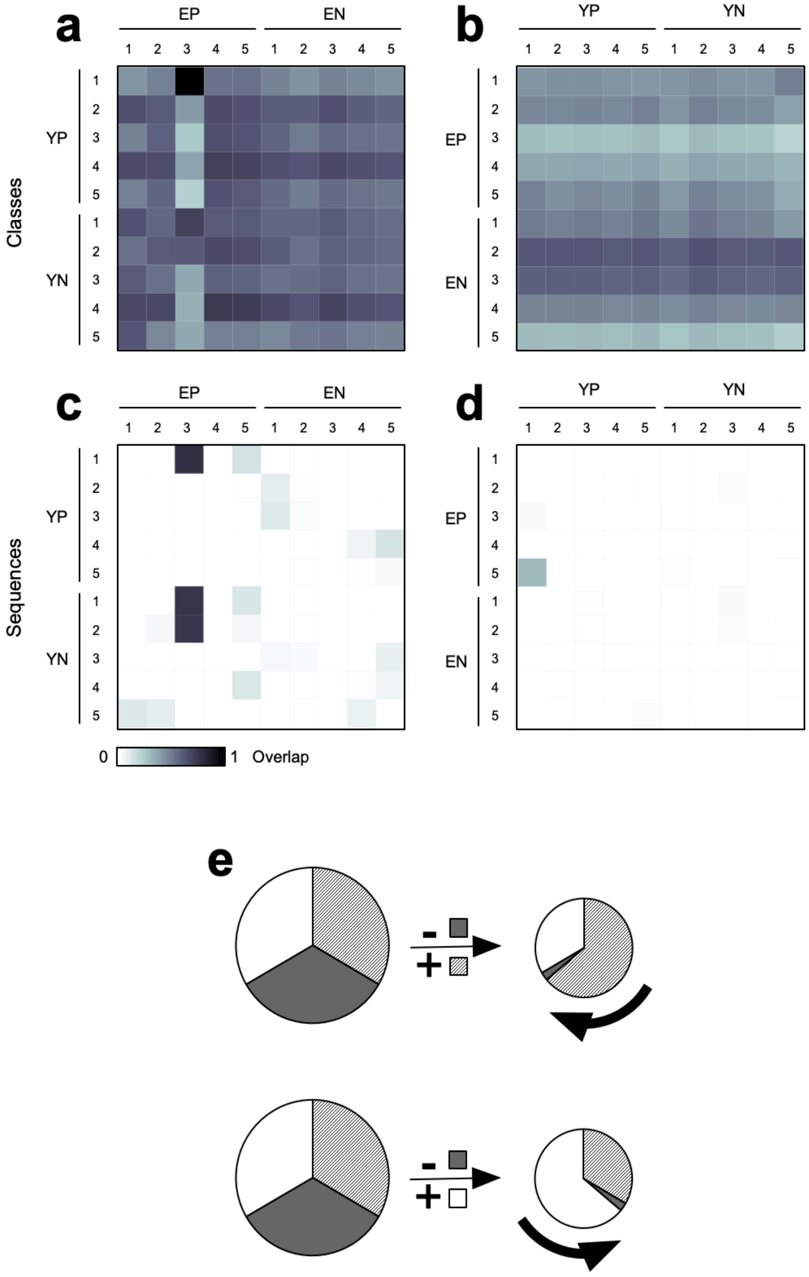

a
EP
EN
1 2 3 4 5 1 2 3 4 5
YP
YN
Classes
b
YP
YN
EP
EN
c
Sequences
d
0
1 Overlap
e
-
+

**Figure S2**

60

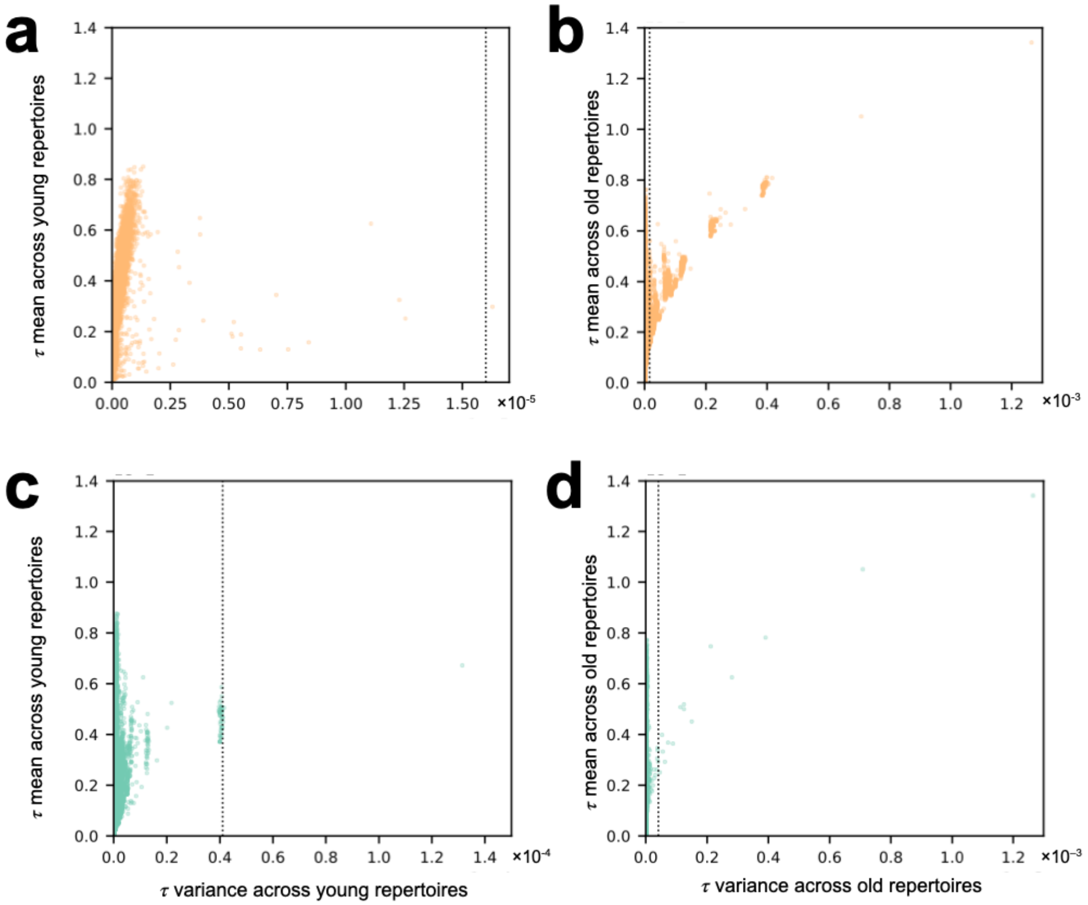
a
b
c
d
τ mean across young repertoires
τ mean across old repertoires
τ variance across young repertoires
τ variance across old repertoires
×10⁻⁵
×10⁻³
×10⁻⁴
×10⁻³
1.4
1.2
1.0
0.8
0.6
0.4
0.2
0.0
0.00 0.25 0.50 0.75 1.00 1.25 1.50
0.0 0.2 0.4 0.6 0.8 1.0 1.2
0.0 0.2 0.4 0.6 0.8 1.0 1.2 1.4
